# *Mycoplasma genitalium* incidence, persistence, concordance between partners and progression: systematic review and meta-analysis

**DOI:** 10.1101/400713

**Authors:** Manuel Cina, Lukas Baumann, Dianne Egli-Gany, Florian S Halbeisen, Hammad Ali, Pippa Scott, Nicola Low

**Affiliations:** Institute of Social and Preventive Medicine, University of Bern, Switzerland; Kirby Institute, University of New South Wales, Australia; University of Otago, New Zealand

## Abstract

**Background:** *Mycoplasma genitalium* is increasingly seen as an emerging sexually transmitted pathogen, and has been likened to *Chlamydia trachomatis*, but its natural history is poorly understood. The objectives of this systematic review were to determine *M. genitalium* incidence, persistence, concordance between sexual partners, and the risk of pelvic inflammatory disease (PID).

**Methods:** We searched Medline, EMBASE, LILACS, IndMed and African Index Medicus from 1 January 1981 until 17 March 2018. Two independent researchers screened studies for inclusion and extracted data. We examined results in forest plots, assessed heterogeneity and conducted meta-analysis where appropriate. Risk of bias was assessed for all studies.

**Results:** We screened 4634 records and included 17 studies; five (4100 women) reported on incidence, five (636 women) on persistence, 10 (1346 women and men) on concordance and three (5139 women) on PID. Incidence in women in two very highly developed countries was 1.07 per 100 person-years (95% CI, 0.61 to 1.53, I2 0%). Median persistence of *M. genitalium* was estimated from one to three months in four studies but 15 months in one study. In ten studies measuring *M. genitalium* infection status in couples, 39-50% of male or female sexual partners of infected participants also had *M. genitalium* detected. In prospective studies, the incidence of PID was higher in women with *M. genitalium* than those without (RR 1.68, 95% CI 0.59 to 2.77, I^2^ 0%, 2 studies).

**Discussion:** Based on findings from this and our linked review of prevalence, concordant *M. genitalium* might be less common than for *C. trachomatis* and the age distributions of the infections differ. The synthesised data about prevalence, incidence and persistence of *M. genitalium* infection are inconsistent. Taken together with evidence about antimicrobial resistance in the two infections, *M. genitalium* is not the new chlamydia.

**Registration Numbers:** PROSPERO: CRD42015020420, CRD42015020405

**KEY MESSAGES:**

- There are calls for widespread screening for *Mycoplasma genitalium*, but the natural history of this emerging sexually transmitted pathogen is poorly understood.
- *M. genitalium* incidence was 1.07 (95% confidence intervals, CI 0.61 to 1.53) per 100-person years in women in highly developed countries, 39-50% of infected individuals had a heterosexual partner with *M. genitalium* and the risk ratio for pelvic inflammatory disease was 1.68 (95% CI 0.59 to 2.77).
- The duration of untreated *M. genitalium* infection is probably longer than persistent detection of *M. genitalium*, as measured in most cohort studies, in which inadvertent treatment cannot be ruled out.
- The results of this systematic review and other evidence sources show important differences in the epidemiology and dynamics of *M. genitalium* and *Chlamydia trachomatis* infection.

## INTRODUCTION

*Mycoplasma genitalium* is increasingly seen as an emerging sexually transmitted pathogen.[1-5] *M. genitalium* is a cause of non-gonococcal urethritis[1] and cervicitis[3] and associations with pelvic inflammatory disease (PID), other reproductive tract complications in women and adverse pregnancy outcomes have been found.[3] *M. genitalium* has thus been called the ‘new chlamydia’.[6] *In a previous systematic review, we found a prevalence of M. genitalium* of approximately 1% in sexually active heterosexuals in the general population, which is similar to that reported for *Chlamydia trachomatis* aged 16-44 years.[7] In sex workers, men who have sex with men (MSM) and clinic-based populations prevalence was higher and more variable.[8] The increasing availability of nucleic acid amplification tests (NAAT) that detect *M. genitalium* has resulted in calls for widespread testing of asymptomatic populations.[9] *But increased testing and treatment is likely to exacerbate the already high proportion of antimicrobial resistant M. genitalium*, since *de novo* resistance to macrolides emerges rapidly.[10 11]

Mathematical modelling could help to understand the balance of benefits and harms that widespread testing and treatment interventions bring.[12] To develop mathematical models, we need robust empirical estimates about the dynamics and probability of progression to PID of *M. genitalium* infection. Important variables are in the following equations:

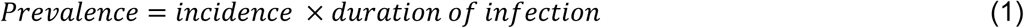

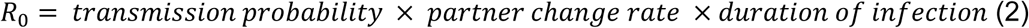

where R_0_ is the basic reproductive number, the average number of secondary infections spread by an infected individual; if R_0_ is greater than one, a pathogen can spread in the population.[13] These measures cannot all be observed directly,[14] but concordant *M. genitalium* status between sexual partners can be used to derive the transmission probability[15] and the duration of persistence of untreated *M. genitalium* can be used to derive the duration of infectiousness.[16] Smieszek and White observed inconsistencies in selected estimates of the clearance rate of *M. genitalium* infection,[17] but did not review these studies systematically. The objectives of this study were to systematically review the research literature to estimate: the incidence of *M. genitalium* infection, persistence of untreated *M. genitalium*, concordance of *M. genitalium* detection, and the risk of developing PID.

## METHODS

This systematic review is one of two linked reviews that used a single search strategy and are described in two protocols.[18 19] A review of the prevalence of *M. genitalium* has been published.[8] We report our findings according to the Preferred Reporting Items for Systematic Reviews and Meta-Analyses (PRISMA, online supplementary file 1).[20]

### Eligibility criteria

We included studies of *M. genitalium* detected by NAAT. Study populations were women and men older than 13 years of age in any country. Eligible study designs were: for incidence of *M. genitalium*, cohort studies with participants who were uninfected at baseline; for persistence, cohort studies that followed people with untreated *M. genitalium* infection; for concordance, cross-sectional studies which enrolled couples or sexual partners of index cases; for incidence of PID, cohort or nested case-control studies that compared women with and without *M. genitalium*.

### Information sources and search strategy

We searched Medline and EMBASE databases, with no language restrictions from 1 January 1981 until 12 July 2016 and updated the search to 17 March 2018. We used thesaurus headings and free text terms that combined *Mycoplasma* and *Mycoplasma genitalium* with genital tract complications (online supplementary file 2, Text S1). We also searched the African Index Medicus, IndMED and LILACS, using the term *Mycoplasma genitalium*. Records were managed using EndNote (version X8.1, Clarivate Analytics, Philadelphia, PA, USA).

### Study selection

Abstracts published before 1 January 1991 were excluded. Two reviewers (MC, LB) assessed study eligibility independently, using a pre-piloted screening form. We resolved differences by discussion or adjudication by a third reviewer (NL).

### Data collection

Two reviewers (MC, LB, DE-G, HA) extracted data independently. Differences were resolved by discussion or adjudication. We extracted data using a standardised, piloted data extraction form in a Research Electronic Data Capture database (REDCap, Vanderbilt University, Nashville, TN, USA). For each study, we extracted study identifiers, topic, language, funding sources, ethical committee approval, use of informed consent, links to other publications, methods, baseline characteristics and study results. For studies reporting persistence of *M. genitalium* only in graphs, we used Plot Digitizer software[21] to record numerical data. We labelled studies with the country in which the data were collected and added consecutive numbers for studies subsequently identified from the same country. Studies reported in the linked systematic review of *M. genitalium* prevalence have the same study identifier (Table S1).[8]

### Risk of bias in individual studies

For cohort or nested case-control studies, we adapted a tool published by the Cochrane Bias Methods Group.[22] For cross-sectional studies, we applied a previously used checklist.[8 23]

### Summary measures

We defined incidence in cohort studies as the rate of new *M. genitalium* infections per 100 person-years of observation in individuals with a negative *M. genitalium* test, either at baseline or a negative test of cure following treatment of a prevalent infection. We defined persistence of *M. genitalium* infection in cohort studies as the proportion of study participants at each follow-up visit with a positive test result. We assessed concordance of *M. genitalium* infection status in cross-sectional studies as the proportion of sexual partners of an infected index case that had a positive test result. We assessed the development of PID in cohort studies and calculated the odds ratio or risk ratio for PID in participants with and without *M. genitalium* infection at baseline.

### Synthesis of results

We used Stata (version 13.1, StataCorp, College Station, TX, USA) for statistical analysis. We examined data about incidence, concordance and PID in forest plots. We stratified studies reporting incidence according to the level of development of the country in which the study was conducted, categorised as very high, high, medium and low using the United Nations Development Programme Human Development Index (HDI).[24] We stratified studies reporting concordance according to study design: studies can enrol couples irrespective of infection status and test all individuals for *M. genitalium* (referred to as partner studies), or can enrol an index case with *M. genitalium* and then test their partners (referred to as index case studies). We calculated the percentage concordance (with 95% confidence intervals, CI) separately for women and men. We assessed the percentage of study variability between studies caused by heterogeneity other than that due to chance with the I2 statistic.[25] Meta-analysis was conducted when deemed appropriate using fixed or random effects models. For estimates of incidence, we estimated a summary estimate of the incidence rate per 100 person years of follow-up (with 95% CI). For concordance, we applied the Freeman-Tukey arcsine transformation to the proportions before meta-analysis and back transformed the summary estimate and its 95% CI.

The data about persistence of *M. genitalium* are presented graphically (Excel:mac 2008, version 12.3.6, Microsoft Corporation, Redmond, WA, USA) without statistical analysis because we anticipated that the results would be too heterogeneous to combine.[17] We conducted a subgroup analysis, using a test of interaction, of differences in concordance by study design.

### Risk of bias across studies

We did not test for small study biases with funnel plots because of the small number of included studies.

## RESULTS

We identified 4634 records and, after exclusion of duplicates and articles published before 1991, we screened 3820 records. We included 17 studies, some of which reported on more than one review question (Table 1, Figure S1). Five studies reported on incidence,[6 26-29] five reported on persistence,[6 27 28 30 31] 10 reported on concordance between partners,[30 32-40] and three studies reported on development of PID.[6 41 42]

**Table 1.** Included studies (n=17), ordered according to outcomes reported

| **Study identifier** | **Study population** | **Study design** | **Review topics** |
| --- | --- | --- | --- |
| Great Britain 2[6] | Female students aged ≤27 years; universities and further education colleges, London, Great Britain | Cohort study | Incidence, persistence, PID |
| Kenya 2[27] | Female sex workers aged 18-35 years; Kariobangi Nairobi City Council, Nairobi, Kenya | Cohort study | Incidence, persistence |
| Kenya 3[31] | Female sex workers, median age 35.3 years; municipal STI clinic Mombasa, Kenya | Cohort study | Incidence, persistence |
| Uganda 1[28] | Female sex workers aged 18-40 years; red light areas within southern Kampala, Uganda | Cohort study | Incidence, persistence |
| Australia 3[26] | Young women aged 16-25 years; primary health clinics in Melbourne, Australia | Cohort study | Incidence |

| Study identifier | Study population | Study design | Review topics |
| --- | --- | --- | --- |
| USA 7[30] | Women aged 14-17 years and their partners; urban primary health care centers, Indianapolis, USA | Cohort study, cross-sectional sampling of couples | Persistence, concordance |
| Great Britain 8[34] | Women and their partners; STI clinic, St. Mary's Hospital, London, Great Britain | Cross-sectional | Concordance |
| Great Britain 9[33] | Men and their partners; STI clinic, St. Mary's Hospital, London, Great Britain | Cross-sectional | Concordance |
| Peru 1[35] | Couples, men aged 19-60 years, women aged 18-55 years; two STI clinics, Lima, Peru | Cross-sectional | Concordance |
| USA 8[32] | Mexican American and African American women with non-viral STI aged 14-45 years and their male partners; San Antonio Metropolitan Health District STI Clinic, USA | Cross-sectional | Concordance |
| Sweden 2[36] | Men aged 16-67 years and their partners; Örebro University Hospital STI clinic, Sweden | Index cases and sexual partners | Concordance |
| Sweden 5[37] | Women aged 14-55 years and men aged 17-67 years and their partners; STI clinic, Falun, Sweden | Index cases and sexual partners | Concordance |
| Sweden 11[38] | Women aged 15-54 years and their partners; Örebro University Hospital STI clinic, Sweden | Index cases and sexual partners | Concordance |
| Sweden 12[39] | Male patients with symptomatic recurrent and/or persistent urethritis aged 20-47 years and their partners; STI clinic, Karolinska Hospital, Stockholm, Sweden | Index cases and sexual partners | Concordance |
| Australia 6[40] | Partners of index cases with *M. genitalium*, median age of female heterosexual partners 26 years; male heterosexual partners 28 years; MSM partners 29 years; Melbourne Sexual Health Centre, Australia | Index cases and sexual partners | Concordance |
| Sweden 10[42] | Women after medical or surgical termination of pregnancy aged 17-40 years; gynaecological outpatient department Malmö University Hospital, Sweden | Nested case-control study | PID |
| USA 6[41] | Women after treatment and cure of PID aged 14-37 years; multiple clinical sites in the United States of America | Cohort study | PID |

Abbreviations: NGU, non-gonococcal urethritis; PID, pelvic inflammatory disease; STI, sexually transmitted infection

### Incidence

We included five studies (Table S2),[6 26-29] with a total of 4100 female participants at baseline and follow-up of 3381.5 person-years. Two studies were conducted in countries with a very high HDI, in students (Great Britain 2)[6] and in attendees of primary health care clinics (Australia 3).[26] All three studies from countries with a low HDI were conducted in female sex workers in (Uganda 1, Kenya 2 and Kenya 3).[27 28 31] We excluded one study post-hoc; Balkus et al. reported combined incidence in women in the USA and Kenya,[43] but the results of the Kenyan cohort were described more extensively in the Kenya 3 study.[29] All women in the Uganda 1 study had a positive test for *M. genitalium* at baseline. Incidence was defined as a positive test result in women who had a preceding negative test result.[28] All studies were at risk of bias (Table S3). All studies reported more than 20% loss to follow-up or did not report it.[6 26-29] Only one (Great Britain 2) compared participants followed up until the end of the studies and participants lost to follow-up.[6]

Figure 1 shows that, in countries with a very high HDI, the pooled estimate of incidence was 1.07 per 100 person-years (95% CI 0.61 to1.53, 2 studies, I^2^ 0%).[6 26] The incidence rates in studies conducted among female sex workers were higher and too heterogeneous to combine (I^2^ 96.7%).

**Figure 1.**
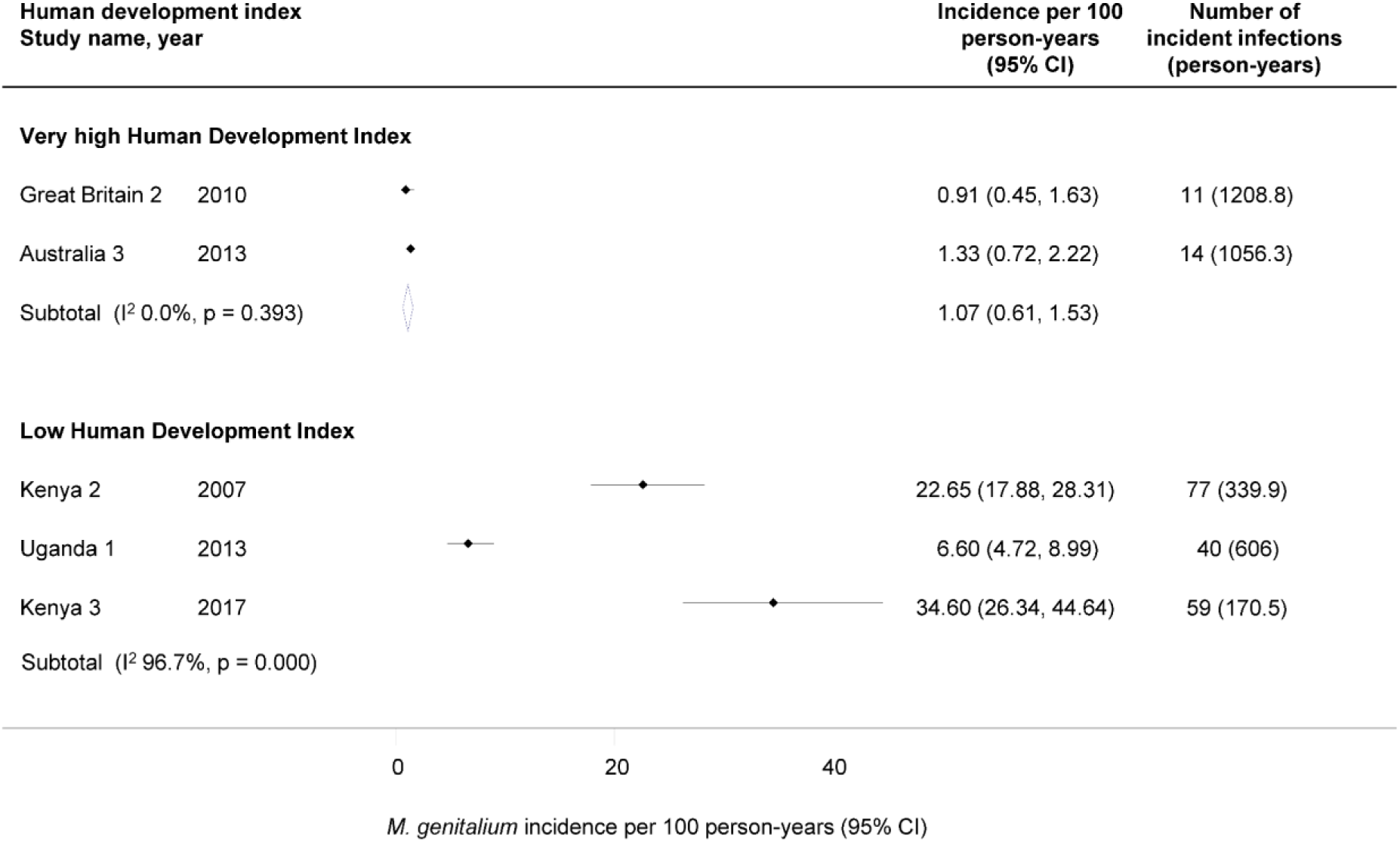
Incident *M. genitalium* infections per 100 person-years by Human Development Index. Solid diamond and lines show the point estimate and 95% CI for each study. The opendiamond shows the point estimate and 95% CI of the summary estimate. The incidence estimates are plotted on a linear scale.

### Persistent detection of *M. genitalium*

We included five publications,[6 27 28 30 31] with a total of 636 female participants at baseline (Table S4). Three studies were conducted in female sex workers in Kenya and Uganda (Uganda 1, Kenya 2 and Kenya 3).[27 28 31] The other two studies were conducted in adolescents enrolled from primary health care facilities (USA 7)[30] and students from educational colleges (Great Britain 2).[6] Duration of follow-up ranged from 12 weeks in USA 7[30] to 33 months in Kenya 2.[27] Specific treatment for *M. genitalium* was not prescribed in any of the studies. All studies were at risk of bias in outcome assessment (Table S5). In Great Britain 2, women with a positive test result for *C. trachomatis* at baseline received antibiotics if they were in the intervention arm of the underlying randomised control trial but could have been treated before the 12-month follow up. In all other studies, participants received either syndromic treatment or treatment for diagnosed *C. trachomatis, N. gonorrhoeae* and/or *Trichomonas vaginalis* at one to three month intervals. Only one study (Great Britain 2) distinguished persistent from re-infections with genotyping.[6] Figure S2 shows a rapid decrease in the proportion of women infected in four studies. Median persistence in the three studies of sex workers was one to three months. The Great Britain 2 study only assessed *M. genitalium* persistence at one subsequent time point at which 25.9% of participants were still infected after a median of 16 months.[6] In USA 7, 31.3% of women remained positive at eight weeks.[30]

### Concordance

We included ten cross-sectional studies,[30 32-39] all of which were conducted in health care facilities (Table S6). Five partner studies enrolled a total of 869 couples irrespective of infection status[30 32-35 40] and five index case studies[36-40] enrolled a total of 477 people with *M. genitalium* and 480 sexual partners. Only the Australia 6 study enrolled MSM.[40] All studies were at risk of bias (Table S7).[30 32-39] The response rate at baseline was only assessed in two studies in which it was below 70% (Great Britain 8 and Peru 1).[33 35] In five of ten studies no standardised procedure to test for *M. genitalium* was reported (Australia 6, Peru 1, Sweden 5, USA 7 and 8).[30 32 35 37 40]

Figure 2 shows overall concordance rates of 39-40% among male partners of women with *M. genitalium* and 40-50% in female partners of infected men, with no marked differences according to study design (Table S8). Concordance among MSM (Australia 6) was 27% (95% CI 19-36%).[40]

**Figure 2.**
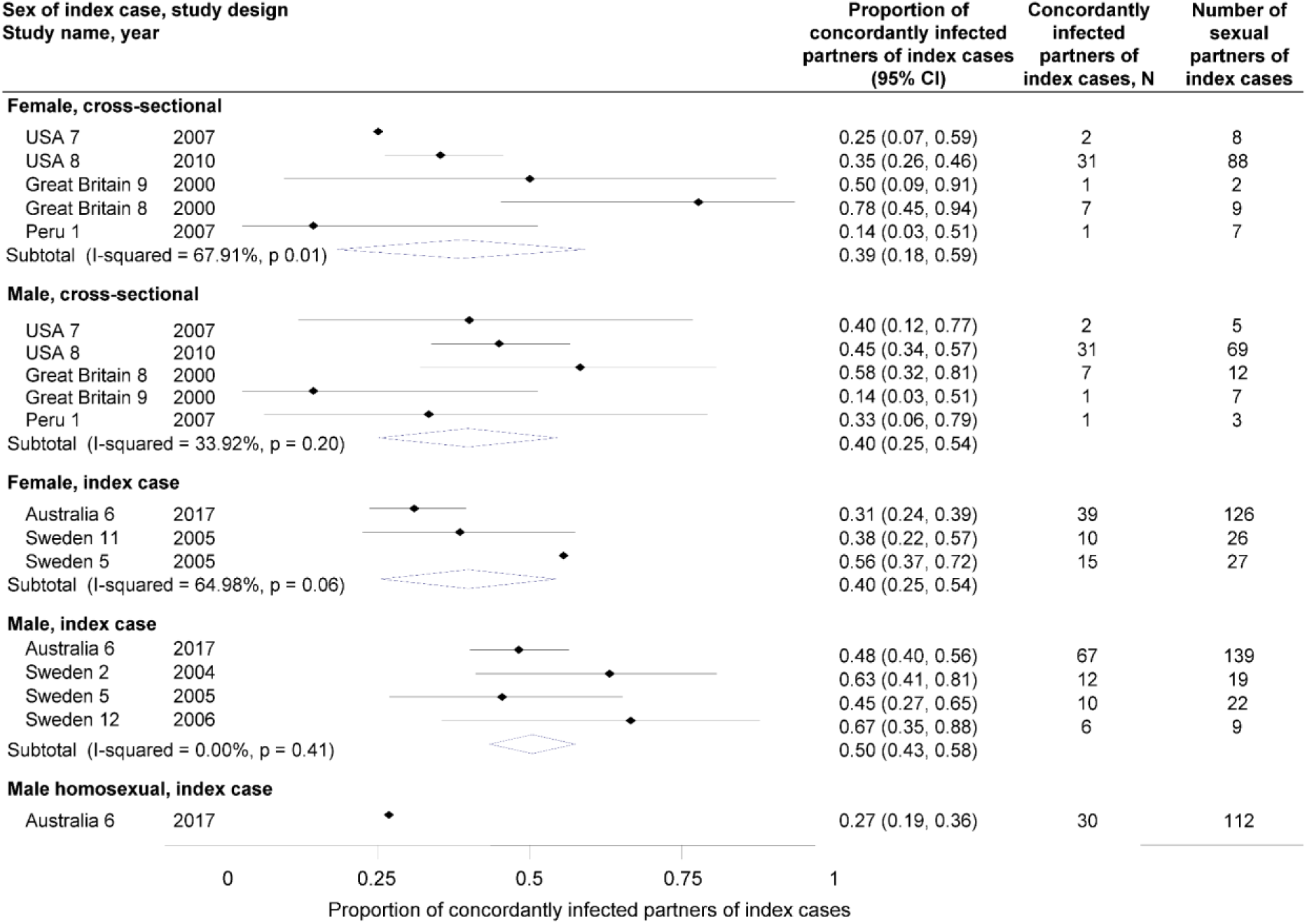
Proportion of concordantly infected sexual partners of individuals with *M. genitalium*, by sex of index case and study design. Solid diamonds and lines show the point estimate and 95% CIs for each study. The open diamond shows the point estimate and the 95% CI of the summary estimate. The proportions are plotted on a linear scale.

### Pelvic inflammatory disease

We included three prospective studies that examined the risk for PID in *M. genitalium*-infected compared with non-infected participants, with a total of 5139 participants at baseline (Table S9).[6 41 42] The Great Britain 2 study enrolled female students in London,[6] USA 6 measured PID recurrence after treatment and cure of a first episode[41] and Sweden 10 was a nested case-control study in women who had undergone medical or surgical termination of pregnancy.[42] PID was diagnosed by endometrial biopsy in USA 6, and using clinical criteria in Great Britain 2[6] and Sweden 10.[41] Follow-up was six weeks in Sweden 10,[42] 12 months in Great Britain 2[6] and 84 months, in USA 6.[41] All studies were at risk of bias (Table S10).[6 41 42] None of the studies assessed if prognostic factors were similar between groups or compared individuals followed up with those lost to follow-up.

All studies found an association between *M. genitalium* and PID (Figure 3). The pooled risk ratio in the two cohort studies was 1.68 (95% CI 0.59 to 2.77, I^2^ 0%). The odds ratio for post-abortion upper genital tract infection was 6.29 (95% CI 1.56 to 25.20).[42]

**Figure 3.**
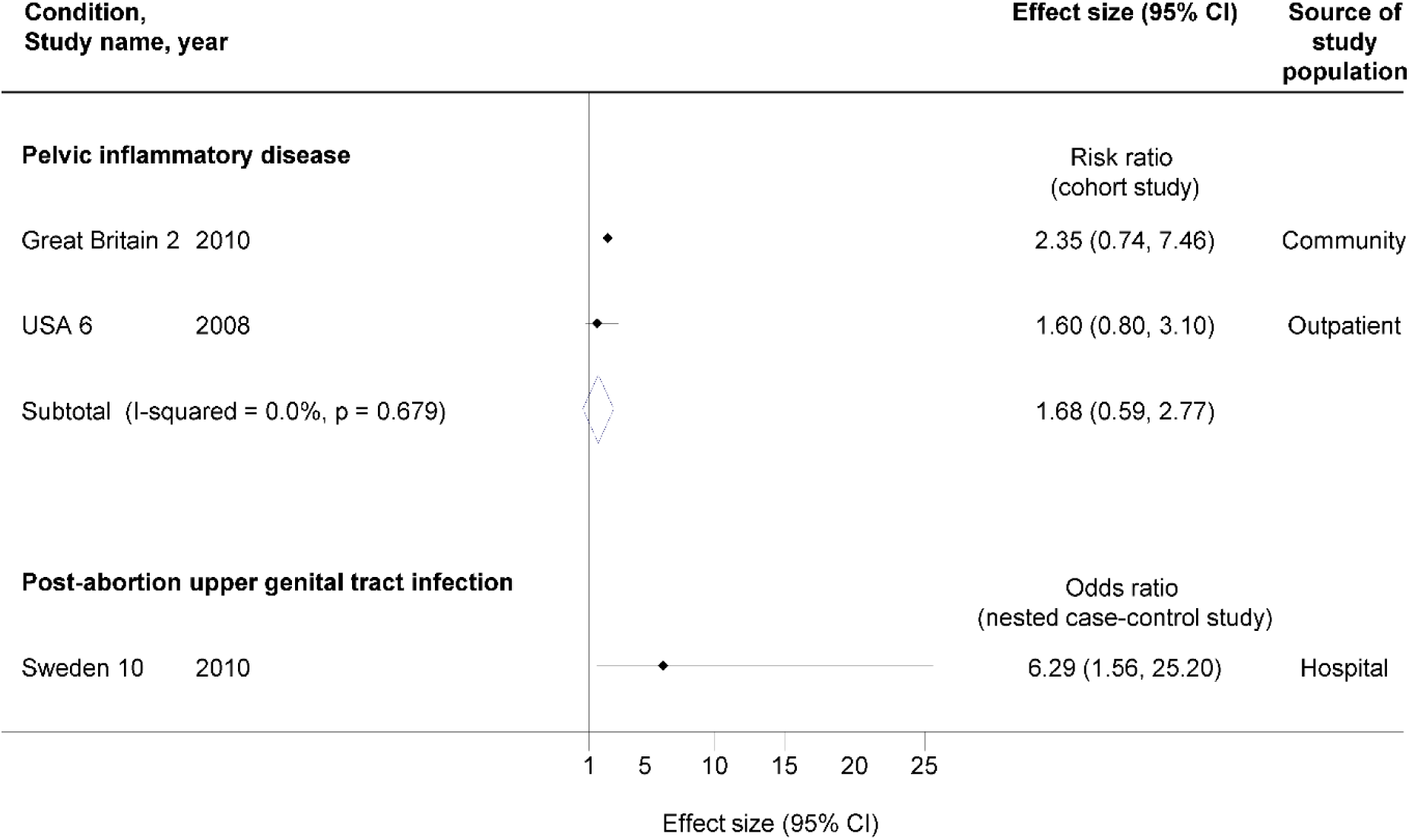
Risk of progression to upper genital tract infection in women with *M. genitalium* compared with women without *M. genitalium*. Solid diamonds and lines show the point estimate and 95% CIs for each study. The open diamond shows the point estimate and the 95% CI of the summary estimate. The effect estimates are plotted on a linear scale.

## DISCUSSION

### Main findings

In this systematic review, the incidence of *M. genitalium* was 1.07 per 100 person-years (95% CI 0.61 to 1.53, I^2^ 0%, 2 studies) in women in very highly developed countries. Median duration of persistence of *M. genitalium* was one to three months in four studies but 15 months in one study. In ten studies measuring *M. genitalium* infection status in heterosexual couples, proportions of concordant results were 39 to 50%. In two prospective studies, the incidence of PID was higher in women with *M. genitalium* than those without (RR 1.68, 95% CI 0.59 to 2.77, I^2^ 0%).

### Strengths and limitations

A strength of our systematic review is the broad search strategy that covered differing topics, which makes it unlikely that we missed important relevant articles. In addition, selection of studies and extraction of data by independent reviewers reduces the risk of errors in data extraction. We assessed the risk of bias in all included studies. The relative importance of the domains of bias affect interpretation depend on the topic. For example, when measuring the duration of persistent detection, accurate assessment of the outcome, untreated infection is important, but most studies were at high risk of bias. The main limitations of the review findings result from the small number of studies overall and between study heterogeneity.

### Interpretation of the findings

The findings of this review and our review of *M. genitalium* prevalence,[8] gives a systematic overview of several important aspects *M. genitalium* infection but reveals inconsistencies in the synthesised evidence about the prevalence, incidence and persistent detection of *M. genitalium* infection (Table S11). Based on equation 1, the quotient of prevalence divided by incidence should be comparable for all studies, but the value obtained is more than one year for the Australia 3,[26] Great Britain 2[6] and Uganda 1[28] studies, and less than one year for Kenya 2[27] and Kenya 3[31] studies. For a sexually transmitted infection that is not highly transmissible to be able to persist (R_0_>1, equation 2) at a low prevalence in the general population, the duration of infectiousness has to be long. The prevalence of *M. genitalium* in the general population of very highly developed countries is around 1%.[8] Concordance between sexual partners of 50% or lower (Figure 3) suggest a low transmission probability. These characteristics are compatible with a longer, rather than a shorter, duration of infectiousness. In the Great Britain 2 and Uganda 1 studies, baseline prevalence was higher than incidence, resulting in an estimated duration of infection of more than one year (Table S11). The rationale and modelling analysis of Smieszek and White favoured a duration of infection similar to the Great Britain 2 study and suggested additional reasons for compatibility with the Uganda 1 study.[17] In all studies, women had opportunities for treatment with antibiotics with some activity against *M. genitalium* at frequencies of as little as a month. The duration of persistent detection was short in all studies that offered treatment every three months or more frequently. With probable inadvertent treatment and re-infection, these cohort studies did not measure the persistence of untreated infection. The uncertainty about the duration of infectiousness of *M. genitalium* contrasts with *C. trachomatis*, for which the literature is extensive and there is broad agreement that prevalence in general populations in high income countries is around 3-4%,[23] incidence is around 4%[44 45] and average duration of infectiousness is slightly more than one year.[46]

The systematic review data suggest some possible differences between *M. genitalium* and *C. trachomatis.* Concordant *M. genitalium* status can be used to estimate the transmission probability of sexually transmitted pathogens.[14 15] Cross-sectional studies of randomly sampled couples, irrespective of infection status, provide the least biased estimate.[14] For this reason, we examined concordance separately in partner studies and in index case studies, but actually found similar estimates in both study designs. In cross-sectional studies, *M. genitalium* concordance was 39-40%. In comparison, *C. trachomatis* concordance in a large cross-sectional study in the USA was 68% (95% CI 56, 78%) for male partners and 70% (58, 80%) for female partners.[47] Findings from our systematic review of *M. genitalium* prevalence suggested that, whilst overall population prevalence of the two infections is similar, *C. trachomatis* positivity is concentrated in younger age groups.[23] M. genitalium was associated with PID in prospective studies (RR 1.68, 95% CI 0.59 to 2.77), but the wide confidence intervals were compatible with no association. This point estimate was lower than that found in by Lis et al., but their inclusion of cross-sectional studies and studies of post-abortal PID in the same meta-analysis might have overestimated the association.[3] The increase in risk of PID following *C. trachomatis* is around 1.8 to 2.8.[48 49] Using data from the Great Britain 2 study and taking into account the low population prevalence of *M. genitalium*, Oakeshott et al. estimated that the population attributable fraction of PID due to *M. genitalium* was about 4%.[6]

### Implications for research and practice

This review adds to the evidence about the biology, dynamics and natural history of *M. genitalium* as a sexually transmitted pathogen. Additional empirical research is needed to provide robust data about the epidemiology of *M. genitalium* infection in men, and to determine the persistence of untreated *M. genitalium*. In the context of evidence of a high probability of emergence of *de novo* resistance and high prevalence of macrolide resistance in *M. genitalium*,[10 11] which do not affect *C. trachomatis*, measures for the management and control of these infections are likely to differ. The estimates from this systematic review can be used in mathematical modelling studies to investigate inconsistencies in the data, to elucidate differences between the transmission dynamics of *M. genitalium* and *C. trachomatis* and to investigate the potential benefits and harms of control interventions. Based on findings from this review, our linked review of prevalence,[8] and evidence about antimicrobial resistance, *M. genitalium* is not the new chlamydia.

## Contributors

Conceived and designed the review: LB, MC, NL, PS. Screened titles, abstracts and full texts: LB, MC, NL. Extracted the data: HA, LB, MC, DE-G. Analysed the data: LB, MC, FH. Wrote the first draft: MC. Revised the paper before submission: HA, LB, MC, DE-G, FH, NL, PS. Approved the final version: HA, LB, MC, DE-G, FH, NL, PS.

## Acknowledgements

We would like to thank Myrofora Goutaki, who gave advice during the review process, Gian-Reto Lohrer, who assisted with data extraction and Dominque Cadosch, who helped with the analysis of persistence data.

## References

1. Taylor-Robinson D, Jensen JS. Mycoplasma genitalium: from Chrysalis to multicolored butterfly. Clin Microbiol Rev 2011;24(3):498–514.

2. Sethi S, Singh G, Samanta P, et al. Mycoplasma genitalium: An emerging sexually transmitted pathogen. Ind J Med Res, Supplement 2012;136(DEC):942-55.

3. Lis R, Rowhani-Rahbar A, Manhart LE. Mycoplasma genitalium Infection and Female Reproductive Tract Disease: A Meta-analysis. Clin Infect Dis 2015;61(3):418–26.

4. Cazanave C, Manhart LE, Bebear C. Mycoplasma genitalium, an emerging sexually transmitted pathogen. Med Mal Infect 2012;42(9):381–92.

5. Bernstein K, Bowen VB, Kim CR, et al. Re-emerging and newly recognized sexually transmitted infections: Can prior experiences shed light on future identification and control? PLoS Med 2017;14(12):e1002474.

6. Oakeshott P, Aghaizu A, Hay P, et al. Is Mycoplasma genitalium in women the New Chlamydia? A community-based prospective cohort study. Clin Infect Dis 2010;51(10):1160–6.

7. Sonnenberg P, Ison CA, Clifton S, et al. Epidemiology of Mycoplasma genitalium in British men and women aged 16-44 years: Evidence from the third National Survey of Sexual Attitudes and Lifestyles (Natsal-3). Int J Epidemiol 2015;44(6):1982–94.

8. Baumann L, Cina M, Egli-Gany D, et al. Prevalence of Mycoplasma genitalium in different population groups: systematic review and meta-analysis. Sex Transm Infect 2018

9. Ross JD, Brown L, Saunders P, et al. Mycoplasma genitalium in asymptomatic patients: implications for screening. Sex Transm Infect 2009;85(6):436–7.

10. Cadosch R, Garcia V, Althaus CL, et al. De novo mutations drive the spread of macrolide resistant Mycoplasma genitalium: a mathematical modelling study. bioRxiv 2018:321216.

11. Horner P, Ingle SM, Garrett F, et al. Which azithromycin regimen should be used for treating Mycoplasma genitalium? A meta-analysis. Sex Transm Infect 2018;94(1):14–20.

12. Birger R, Saunders J, Estcourt C, et al. Should we screen for the sexually-transmitted infection Mycoplasma genitalium? Evidence synthesis using a transmission-dynamic model. Sci Rep 2017;7(1):16162.

13. Anderson RM, May RM. Infectious diseases of humans: dynamics and control. Oxford, United Kingdom: Oxford University Press 1991.

14. Garnett GP, Bowden FJ. Epidemiology and control and curable sexually transmitted diseases: opportunities and problems. Sex Transm Dis 2000;27(10):588–99.

15. Althaus CL, Heijne JC, Low N. Towards More Robust Estimates of the Transmissibility of Chlamydia trachomatis Sex Transm Dis 2012;39(5):402–04.

16. Althaus CL, Heijne JCM, Roellin A, et al. Transmission dynamics of Chlamydia trachomatis affect the impact of screening programmes. Epidemics 2010;2(3):123–31.

17. Smieszek T, White PJ. Apparently-different clearance rates from cohort studies of Mycoplasma genitalium are consistent after accounting for incidence of infection, recurrent infection, and study design. PLoS ONE 2016;11(2)

18. Low N, Baumann L, Cina M, et al. Mycoplasma genitalium infection: prognosis and transmissibility [CRD42015020405]. <https://www.crd.york.ac.uk/prospero/display_record.php?RecordID=20405>, (accessed 16 July 2018.).

19. Low N, Cina M, Baumann B, et al. Mycoplasma genitalium infection: prevalence, incidence and persistence [CRD42015020420]. <https://www.crd.york.ac.uk/prospero/display_record.php?RecordID=20420>, (accessed 16 July 2018.).

20. Liberati A, Altman DG, Tetzlaff J, et al. The PRISMA statement for reporting systematic reviews and meta-analyses of studies that evaluate health care interventions: explanation and elaboration. Ann Intern Med 2009;151(4):W65–94.

21. Howaldt J. Plot Digitizer. <http://plotdigitizer.sourceforge.net/>, (accessed 19 June 2017.).

22. Cochrane Collaboration. Tool to Asseess Risk of Bias in Cohort Studies. <https://methods.cochrane.org/bias/sites/methods.cochrane.org.bias/files/public/uploads/Tool%20to%20Assess%20Risk%20of%20Bias%20in%20Cohort%20Studies.pdf>, (accessed 15 August 2018.).

23. Redmond SM, Alexander-Kisslig K, Woodhall SC, et al. Genital Chlamydia prevalence in europe and non-European high income countries: systematic review and meta-analysis. PLoS One 2015;10(1):e0115753.

24. United Nations Development Programme. Human Development Report 2014: Sustaining Human Progress-Reducing Vulnerabilities and Building Resilience. New York, USA, 2014.

25. Higgins JP, Thompson SG. Quantifying heterogeneity in a meta-analysis. Stat Med 2002;21(11):1539–58.

26. Walker J, Fairley CK, Bradshaw CS, et al. Mycoplasma genitalium incidence, organism load, and treatment failure in a cohort of young Australian women. Clin Infect Dis 2013;56(8):1094–100.

27. Cohen CR, Nosek M, Meier A, et al. Mycoplasma genitalium infection and persistence in a cohort of female sex workers in Nairobi, Kenya. Sex Transm Dis 2007;34(5):274–9.

28. Vandepitte J, Weiss HA, Kyakuwa N, et al. Natural history of Mycoplasma genitalium infection in a cohort of female sex workers in Kampala, Uganda. Sex Transm Dis 2013;40(5):422–7.

29. Balkus JE, Manhart LE, Lee J, et al. Periodic Presumptive Treatment for Vaginal Infections May Reduce the Incidence of Sexually Transmitted Bacterial Infections. J Infect Dis 2016;213(12):1932–37.

30. Tosh AK, Van Der Pol B, Fortenberry JD, et al. Mycoplasma genitalium among adolescent women and their partners. J Adolesc Health 2007;40(5):412–7.

31. Lokken EM, Balkus JE, Kiarie J, et al. Association of Recent Bacterial Vaginosis With Acquisition of Mycoplasma genitalium. Am J Epidemiol 2017;186(2):194–201.

32. Thurman AR, Musatovova O, Perdue S, et al. Mycoplasma genitalium symptoms, concordance and treatment in high-risk sexual dyads. Int J STD AIDS 2010;21(3):177–83.

33. Keane FE, Thomas BJ, Gilroy CB, et al. The association of Chlamydia trachomatis and Mycoplasma genitalium with non-gonococcal urethritis: observations on heterosexual men and their female partners. Int J STD AIDS 2000;11(7):435–9.

34. Keane FE, Thomas BJ, Gilroy CB, et al. The association of Mycoplasma hominis, Ureaplasma urealyticum and Mycoplasma genitalium with bacterial vaginosis: observations on heterosexual women and their male partners. Int J STD AIDS 2000;11(6):356–60.

35. Nelson A, Press N, Bautista CT, et al. Prevalence of sexually transmitted infections and high-risk sexual behaviors in heterosexual couples attending sexually transmitted disease clinics in Peru. Sex Transm Dis 2007;34(6):344–61.

36. Falk L, Fredlund H, Jensen JS. Symptomatic urethritis is more prevalent in men infected with Mycoplasma genitalium than with Chlamydia trachomatis. Sex Transm Infect 2004;80(4):289–93.

37. Anagrius C, Lore B, Jensen JS. Mycoplasma genitalium: prevalence, clinical significance, and transmission. Sex Transm Infect 2005;81(6):458–62.

38. Falk L, Fredlund H, Jensen JS. Signs and symptoms of urethritis and cervicitis among women with or without Mycoplasma genitalium or Chlamydia trachomatis infection. Sex Transm Infect 2005;81(1):73–8.

39. Wikstrom A, Jensen JS. Mycoplasma genitalium: a common cause of persistent urethritis among men treated with doxycycline. Sex Transm Infect 2006;82(4):276–9.

40. Slifirski JB, Vodstrcil LA, Fairley CK, et al. Mycoplasma genitalium Infection in adults reporting sexual contact with infected partners, Australia, 2008-2016. Emerg Infect Dis 2017;23(11):1826–33.

41. Haggerty CL, Totten PA, Astete SG, et al. Failure of cefoxitin and doxycycline to eradicate endometrial Mycoplasma genitalium and the consequence for clinical cure of pelvic inflammatory disease. Sex Transm Infect 2008;84(5):338–42.

42. Bjartling C, Osser S, Persson K. The association between Mycoplasma genitalium and pelvic inflammatory disease after termination of pregnancy. BJOG 2010;117(3):361–4.

43. Balkus JE, Manhart LE, Jensen JO, et al. Detection of macrolide resistance-mediating mutations among women with mycoplasma genitalium infection in the preventing vaginal infections trial. Am J Obstet Gynecol 2016;215(6):S828.

44. Scott Lamontagne D, Baster K, Emmett L, et al. Incidence and reinfection rates of genital chlamydial infection among women aged 16-24 years attending general practice, family planning and genitourinary medicine clinics in England: a prospective cohort study by the Chlamydia Recall Study Advisory Group. Sex Transm Infect 2007;83(4):292–303.

45. Walker J, Tabrizi SN, Fairley CK, et al. Chlamydia trachomatis incidence and re-infection among young women--behavioural and microbiological characteristics. PloS One 2012;7(5):e37778.

46. Davies B, Anderson SJ, Turner KM, et al. How robust are the natural history parameters used in chlamydia transmission dynamic models? A systematic review. Theor Biol Med Model 2014;11:8.

47. Quinn TC, Gaydos C, Shepherd M, et al. Epidemiologic and microbiologic correlates of Chlamydia trachomatis infection in sexual partnerships. JAMA 1996;276(21):1737–42.

48. Reekie J, Donovan B, Guy R, et al. Risk of Pelvic Inflammatory Disease in Relation to Chlamydia and Gonorrhea Testing, Repeat Testing, and Positivity: A Population-Based Cohort Study. Clin Infect Dis 2018;66(3):437–43.

49. Davies B, Turner KM, Leung S, et al. Comparison of the population excess fraction of Chlamydia trachomatis infection on pelvic inflammatory disease at 12-months in the presence and absence of chlamydia testing and treatment: Systematic review and retrospective cohort analysis. PLoS One 2017;12(2):e0171551.

